# Acute ketamine treatment produces long-term anxiolytic effects despite increasing oxidative stress in female Wistar Kyoto rats

**DOI:** 10.64898/2026.06.26.734907

**Authors:** Alena Lemeshova, Feysal Abdirahaman, Haney Haidari, Cherish Zhao, Aiman Limbada, Jennifer A. Honeycutt

## Abstract

Treatment-resistant depression and anxiety remain major challenges in psychiatry, particularly in female patients, who are disproportionately affected yet remain underrepresented in preclinical ketamine research. The present study investigated short- and long-term anxiolytic effects of acute subanesthetic ketamine administration in female Wistar-Kyoto (WKY) rats, a validated genetic model of treatment-resistant affective dysfunction. Subjects received a single intraperitoneal injection of saline vehicle or racemic ketamine (5, 10, or 15 mg/kg), followed by acoustic startle response (ASR) testing 24 hours and 7 days later. Oxidative stress was assessed using 8-oxo-2’-deoxyguanosine (8-oxo-dG) immunofluorescence in the basolateral amygdala (BLA), prefrontal cortex (PFC), and hippocampus, alongside analysis of parvalbumin-positive (PV+) interneurons. Ketamine treatment produced dose- and time-dependent behavioral effects with 10 mg/kg eliciting the strongest delayed anxiolytic-like response at 7 days, while 15 mg/kg showed more immediate behavioral effects at 24 hours. While ketamine did not alter PV+ cell count, it significantly increased oxidative stress markers globally in the BLA and prelimbic region of the PFC and specifically in the PV+ interneurons in the BLA. The findings suggest that ketamine’s therapeutic effects in female WKY rats may involve region-specific modulation of stress circuitry and oxidative signaling rather than gross interneuron loss. Overall, the study provides evidence for sex-dependent and temporally dynamic effects of ketamine in a translational model of treatment-resistant anxiety and depression.

## Introduction

One of the most widely researched topics in neuropharmacology is the management of treatment-resistant disorders, such as anxiety and post-traumatic stress disorder (PTSD) which can be unresponsive to commonly used selective serotonin reuptake inhibitors (SSRIs), tricyclic antidepressants (TCAs), serotonin/norepinephrine reuptake inhibitors (SNRIs), and/or monoamine oxidase inhibitors (MAOIs). Despite the broad range of standard antidepressants available, about 30-60% of anxiety disorder patients continue to experience symptoms after the start of treatment, with 12-20% of individuals experiencing relapse after three years (Bokma et al. 2021). PTSD displays a similar pattern, in which remission rates following classic treatment range from 20-30%, and 40-60% demonstrate resistance traits (Marseille et al. 2020; Almeida et al. 2024). Unlike the commonly prescribed treatments, ketamine, a non-competitive N-methyl-D-aspartic acid (NMDA) receptor antagonist, produces a swift therapeutic response in resistant patients with effects that last as long as seven days after a single administration by directly affecting glutamate modulation (Gass et al. 2019; Murrough et al. 2013; Yang et al. 2016). In their systematic review of studies involving ketamine infusions in treatment-resistant depression (TRD) patients, Serafini et al. (2014) found that ketamine infusions demonstrated robust antidepressant properties in the majority of studies, indicating its potential as an alternative to existing therapies. The effect was reported to occur as rapidly as within one day of a single subanesthetic infusion and last for at least three to fourteen days in studies on MDD patients (Singh et al., 2017; Ponton et al., 2022; Yavi et al., 2022).

Ketamine affects monoaminergic, glutamatergic, muscarinic, and opioidergic systems, although its antidepressant effect is thought to stem from the induced blockade of NMDA receptors, which potentiates synaptogenesis and neuroplasticity by partially reversing gamma aminobutyric acid (GABA)-related dysfunction recorded in multiple affective disorders (Yavi et al., 2022). Despite the drug’s promising effects, the preclinical studies investigating ketamine as a psychiatric treatment lack standardization in experimental design, which raises a challenge in determining the most fitting medically relevant dose in rodents (Lemeshova et al., 2026). In a review paper our laboratory has published in 2026, acute lower doses of 10-30 mg/kg administered peritoneally emerged as most effective, especially when the behavioral measures were conducted 24 hours or more after administration, allowing for disambiguation of acute locomotor and cognitive effects from potential therapeutic benefits (Lemeshova et al., 2026). In a variety of studies on ketamine as an anxiolytic 10 mg/kg dose demonstrates the best sustained amelioration of maladaptive anxious-like behavioral profile, while the dose of 15 mg/kg emerges to be anxiogenic, especially when administered repeatedly (Lemeshova et al., 2026).

The lack of understanding behind ketamine’s rapid effects and susceptibility of patients based on their genetic predisposition to mental health disorders requires an appropriate preclinical model of affective dysfunction. In rats, the baseline antidepressant dose responsiveness is much lower than in human patients (Will et al., 2003), leading to issues with translatability of preclinical work. However, this increased sensitivity is not observed in the inbred strain of Wistar Kyoto (WKY) rats, which exhibit depressive and anxiety-like behaviors comparable to those of human patients (Will et al., 2003; Edwards and King, 2009; McAuley et al. 2009). For instance, Li et al. (2022) recorded increased immobility in the forced swim test (FST; suggestive of learned helplessness; Can et al., 2011), decreased sucrose preference in the sucrose preference test (an indicator of anhedonia; Primo et al., 2023), and less time spent in the center of the open field test (an indicator of increased anxiety; Kraeuter et al. 2018). These animals are not responsive to serotonergic drugs, such as citalopram, fluoxetine, and 8-OH-DPAT (López-Rubalcava, 2000), but demonstrate pronounced behavioral changes following administration of ketamine, such as an increase in swim time and latency to stop swimming in the FST after an i.p. injection of 10 and 30 mg/kg of ketamine (Manduca et al., 2020). Therefore, this preclinical model allows for pharmaceutical treatment testing without the need to subject rats to additional stress, as well as makes for a higher degree of translatability to human patients with treatment resistance (Redei et al. 2023).

Central to affective dysfunction is the prefrontal cortex (PFC)–basolateral amygdala (BLA) stress circuitry, which is modulated by GABAergic, parvalbumin-expressing (PV+) interneurons (Binette et al., 2023). As PV+ interneurons suppress excessive excitatory output and maintain the excitation/inhibition balance, a failure to properly modulate signals between PFC and BLA results in numerous neuropsychiatric conditions, including anxiety, MDD, and PTSD (Iqbal et al., 2023; Bogdańska-Chomczyk et al., 2024; Su et al., 2025). In a study on anxiety-like behavior following chronic stress, Ryazantseva et al. (2025) reported that reduced GABA release from PV+ cells caused hyperexcitability in the amygdala projection neurons, suggesting that PV is crucial for regulation fear and emotional learning and shaping the outputs of PFC and BLA.

PV+ cells are among the primary targets of ketamine action due to their high expression of NMDA 2A receptors, which ketamine inhibits in a sex- and region-specific manner (Perlman et al., 2021). Rat models of schizophrenia induced by chronic administration of ketamine demonstrate a reduction in the number of PV+ cells in the PFC paired with reports of abnormally high increase in the cell number in the CA3 field of hippocampus (Sabbagh et al. 2013). While less information is available about the specifics of PV+ cell expression and number in the WKY strain, its control strain counterpart, the spontaneously hypertensive (SHR) rat line, is reported to demonstrate a reduced PV cell number in prelimbic, anterior cingulate, and primary and secondary motor cortices (Bogdańska-Chomczyk et al., 2024). In their 2012 paper, Jiao et al. demonstrated abnormal activation patterns of PV-positive neurons in the BLA compared to the control Sprague Dawley outbred strain, specifically via less activation following a potent stressor, which could point toward weaker inhibitory regulation of amygdala circuits via PV cells in the strain. Together, these findings suggest that stress-related NMDA-receptor dysfunction in WKY rats may co-occur with PV-interneuron abnormalities like those observed in SHR rats, pointing to a potential PV–NMDA2A interaction contributing to the strain’s behavioral and stress-sensitivity phenotypes.

While monoamine imbalance is the first mechanism to be commonly linked to psychiatric conditions, other processes, such as neuroinflammation, have been a focus of investigation in the context of anxiety, MDD, and other disorders (Yavi et al., 2022). Oxidative stress is a major factor in the inflammation hypothesis, arising from an imbalance of oxidant and antioxidant compounds that results in excessive free radical generation, damaging biomolecules and augmenting the production of pro-inflammatory molecules such as cytokines and chemokines (Réus et al., 2015). Reactive oxygen and nitrogen species, including other free radicals, are balanced by antioxidant defenses, which ketamine potentially aids at subanesthetic doses (Réus et al., 2015). Thus, Liang et al. (2018) found a significant restoration in levels of superoxide dismutase, a crucial antioxidant enzyme, in a mouse model of traumatic brain injury following acute ketamine treatment. A similar enhancement of antioxidant enzyme activity in a rodent model of depression following administration of ketamine was reported by Ivanović et al. (2025). Overall, evidence pointing toward ketamine’s conferred neuroprotection against markers of oxidative damage has been reported reliably in multiple preclinical experimental paradigms.

However, reports of decreased oxidation following ketamine treatment are not always consistent. For instance, multiple studies cite an increase in 8-oxo-2’-deoxyguanosine (8-oxo-dG), a modified DNA base that forms when guanine is oxidized by reactive oxygen species, and loss of PV cells as by-products of the drug’s administration in mice (Schiavone et al., 2019) and rats (Liu et al., 2012). Despite the conflicting findings, the intersections of oxidative outcomes, sex-dependent effects, and characterization of PV-interneuron function remain generally unexplored, specifically within the WKY rat model.

Women are twice as likely to develop MDD, anxiety disorders, and PTSD, but most studies on therapeutic ketamine exclusively use male animals (Kessler et al., 2012; Olff, 2017; Salk et al., 2017; Arjmand et al., 2023). The few currently available publications that did include female animals have suggested the preferential effectiveness of a lower ketamine dose compared to male animals, which reinforces the importance of studying sex differences in preclinical trials (Lemeshova et al., 2026; Yang and Chen, 2024). In a review on existing literature studying ketamine as anxiolytic, our laboratory has found that only 8 out of 35 studies included female subjects, and only one study reported anxiolytic effects in females compared to numerous such reports in male animals, underscoring potential underlying sex-dependent mechanisms that require investigation in both sexes (Lemeshova et al., 2026). Despite potential hypotheses for ketamine and estrogen interactions available, existing literature in the field rarely includes female subjects, complicating interpretation of existing data and potentially concealing its sex-dependent effects (Lemeshova et al., 2026).

The present study utilized the WKY strain as a model of genetic predisposition to depression and anxiety to measure behavior in the ASR paradigm, which is highly translatable due to its simple stimulus-physiological response relationship that mimics the exaggerated degree of startle response present in patients with anxiety, MDD, and PTSD (Morgan et al., 1995; Koch and Schnitzler, 1997; Ray et al., 2008; Vaidyanathan et al. 2013). Similar to humans with affective dysfunction, WKY rats were expected to demonstrate maladaptive startle-related behavior (McAuley et al. 2009; Vaidyanathan et al. 2013), which we hypothesized ketamine treatment would normalize both immediately (24 hours after exposure) and long-term (seven days after exposure) following the administration of a 5, 10, or 15 mg/kg dose. Additionally, this treatment was predicted to decrease oxidation measured via 8-oxo-dG in the prefrontal cortex, basolateral amygdala, and hippocampus in a dose-dependent manner, globally and locally within the PV+ cells.

## Methods

### Subjects

#### Animals

Forty-four female adult Wistar Kyoto rats (Charles River Laboratories, Wilmington, MA) arrived at Bowdoin College on postnatal day (PND) 21 and were housed in pairs with *ad libitum* access to water and rodent chow (Lab Diet 5P00 Prolab RMH 3000) in a vivarium that is controlled for temperature and humidity (72°F and 40-60% relative humidity, respectively) with a 12 hour light cycle (lights on at 0700h and off at 1900h). Ketamine treatment was administered on PND 50, and behavioral testing was conducted on PND 51 and 57 (Figure 1).

**Figure 1.**
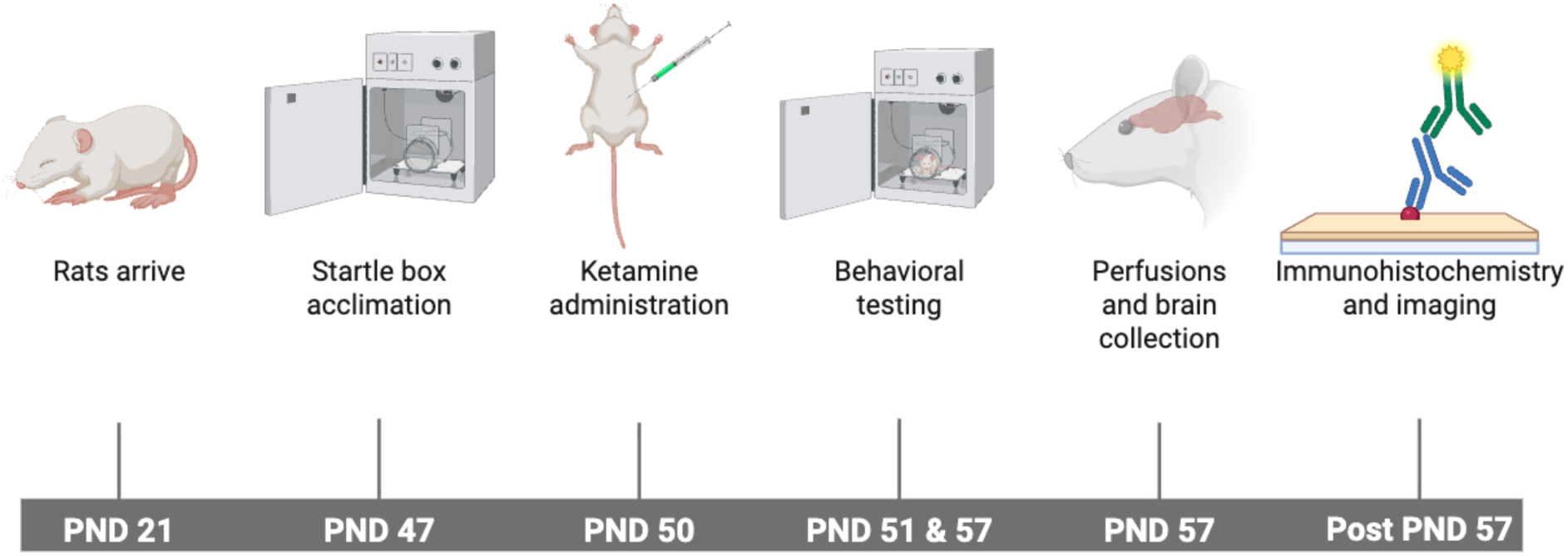
Experimental timeline. Made with BioRender.

#### Experimental Groups

The animals were divided into three experimental treatment groups based on dosage of 5, 10, or 15 mg/kg of ketamine and one control group (4 groups total; n = 10/group). All experiments were performed with approval of Bowdoin College’s Institutional Animal Care and Use Committee in accordance with the Guide for the Care and Use of Laboratory Animals (NIH).

#### Ketamine Treatment

Ketamine hydrochloride stock solution of 100 mg/mL was diluted in 0.9% NaCl saline to inject 5, 10, or 15 mg/kg intraperitoneally (i.p.; Ketaset; Bærentzen et al. 2024; Manduca et al. 2020; McDonnell et al. 2021). For the control group, a comparable volumetric dose of vehicle saline was used. All injections were administered on PND50 during the animals’ light cycle.

### Behavioral Testing

#### Acoustic Startle Response (ASR)

All rats were habituated to the SR-LAB Startle Response System (San Diego Instruments, San Diego, CA) for five minutes each day, for six days with a background noise of 65 dB playing in the inner speakers to reduce possible confounding effects of stress on the expression of the study-relevant neural markers after testing (Palmer and Printz 1999). Upon placement of a rat in the inner chamber of startle box, 65 dB background noise was played for 5 minutes (for acclimation), after which a 100 ms pulse of either 95, 105, or 115-dB white noise followed. Every individual session had 30 randomized trials, each containing 10 pulses of each loudness level to test the behavioral response recorded by the SR-LAB system. The output was collected as average startle, an average value of startle throughout all 30 trials for each subject, and maximum startle, the amplitude of startle response across the 30 trials for each subject. The data was collected twice, during an immediate and a delayed trial, 24 hours and 7 days after the administration of ketamine treatment accordingly.

All original measurements from paw pressure on the floor of the startle chamber during the startle display, recorded in millivolts, were transformed into z-scores to reduce chamber-specific variability and standardize outputs across multiple SR-LAB Startle Response System boxes. Standardization was additionally used to normalize highly variable raw startle amplitudes and facilitate better comparisons across subjects and testing chambers, consistent with methodological recommendations for improving robustness and comparability in acoustic startle response assays (Figure 2; Miller et al., 2020).

**Figure 2.**
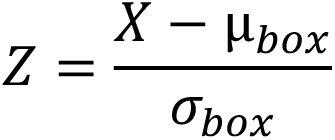
Formula for conversion of startle behavioral data into z-scores: X = recorded response, µ = mean response, σ = standard deviation.

#### Behavioral Data Analysis

Two-way ANOVAs with treatment dose and timing of the test (immediate or delayed) were performed using GraphPad Prism, and Šidák’s post hoc tests were used to assess significant main effects and/or interactions. Cut-off value for statistical significance was set at *p*<0.08 for trends and at *p*<0.05 for statistical significance. Outliers were removed in line with the Grubbs’ outlier test (GraphPad Software).

### Tissue Collection and Processing

#### Brain Extraction

Within an hour of the final acoustic startle response test, animals were euthanized via CO_2_ and transcardially perfused with 0.9% NaCl to remove blood from the tissue followed by 4% paraformaldehyde solution until tissue fixation as judged by the stiffness of the limbs. Rats were then decapitated, and the brains were retrieved and stored in 4% paraformaldehyde solution for a week for postfixing, after which they were transferred to 30% sucrose solution for cryoprotection until fully sunk.

The tissue was then sliced into 40 µm sections with a freezing microtome (Leica Biosystems, Wetzlar, Germany) and stored in well plates with anti-freeze solution (10% 0.2 M phosphate buffer, 30% deionized H2O, 30% ethylene glycol, 30% glycerol) in a -20°C freezer until further analysis. Tissue was saved in serial sections of brain regions involved in depression and anxiety, including prefrontal cortex, basolateral amygdala, and hippocampus (Drevets et al., 2008).

#### Immunohistochemistry for PV and 8-Oxo-dG

The staining protocol to assess immunoreactivity of PV and 8-oxo-dG in the collected tissue was based on previous work (Gürler et al. 2014; Xiao et al. 2017; Gildawie et al. 2020). Sections were washed 3 x 5min with phosphate-buffered saline (PBS), then incubated in RNase solution (2.5 mg/mL RNase A in 5 mM Tris–HCl, 7.5 mM NaCl) for 1 h at 37 °C to prevent anti-8-oxo-dG binding to RNA). The sections were then washed 3 x 5min in PBS and blocked with normal goat serum (NGS, NC9660079, Jackson Immuno) and 0.2% Triton-X in PBS (0.2% PBST). Tissue was washed 3 x 5min and incubated overnight with primary antibodies including: mouse anti-8-oxo-dG (1:300, 4354-MC-050, ThermoFisher) and rabbit anti-PV (1:1000, PA1–933, ThermoFisher) at 4 °C. Next day, sections were washed in PBS and incubated in the following secondary antibodies (raised in goat) for 1 hour at RT: anti-mouse conjugated with Alexa Fluor 568 (1:1000, A11031, ThermoFisher) to visualize 8-oxo-dG and anti-rabbit conjugated with Alexa Fluor 555 (1:1000, A-21428, ThermoFisher) to visualize PV. Sections were washed 3 x 5min in PBS, then mounted on glass slides in a dark room to prevent photobleaching. All washes and incubations were done on an orbital shaker. Slides were dried in a dark drawer and cover-slipped using Prolong Glass Antifade mounting media (Invitrogen).

#### Imaging and Image Analysis

Imaging was done via a Keyence BZ-X800 fluorescent microscope (Osaka, Japan) at 10x magnification for dentate gyrus of the hippocampus, 20x for prefrontal cortex and CA1/CA3 regions of the hippocampus, and 40x for the basolateral amygdala. Two images were taken per brain region (BLA, PL and IL PFC, and CA1, CA3, and DG HPC) for each subject, resulting in 12 images per subject. Cy3 channel was used for PV images at 0.025 s exposure while Cy5 was used for 8-oxo-dG at 1/3.5 s exposure, both on high resolution.

Analysis was conducted using Fiji (ImageJ, NIH), including PV cell count, global 8-oxo-dG intensity, and colocalized PV/8-oxo-dG intensity. For PV cell count, the background was subtracted with a 50.0-pixel rolling ball radius and threshold was set to a brightness of 50-55. Particle analysis was then conducted with a size range of 250-infinity and circularity 0.1 to 1.0. For 8-oxo-dG global intensity, the images were split into blue and red channels, then integrated density values were measured. Colocalization of PV and 8-oxo-dG was achieved using the Merge Channel function for blue and red channels, then the PV cells were hand-traced, and their integrated density was measured for the corresponding areas of the red channel.

#### Imaging Data Analysis

Global 8-oxo-dG intensity, PV cell number, and 8-oxo-dG/PV signal colocalization were analyzed via one-way ANOVAs across the control and experimental conditions using GraphPad Prism, with Tukey’s post hoc tests for pair-wise comparisons. Cut-off value for statistical significance was set at *p*<0.08 for trends and at *p*<0.05 for statistical significance. Outliers were removed in line with the Grubbs’ outlier test (GraphPad Software).

## Results

### Acoustic Startle Response

#### Normalized Average Startle

Following i.p. injections of three ketamine doses, the startle responses were compared 24 hours and 7 days post-administration via a two-way repeated measures ANOVA (time x treatment condition) to determine whether ketamine dose influenced changes in behavior. The analysis revealed a significant interaction between trial timing and treatment (*F*(3,40)=3.183, *p*=0.0341), indicating the effect of ketamine doses was variable depending on whether 24 hours or 7 days had passed since administration (Figure 3). A significant main effect of time was also found (*F*(1,40)=7.735, *p*=0.0082), demonstrating a shift in behavioral scores between the 24-hour and 7-day assessments regardless of treatment. In comparison, treatment alone did not yield a significant main effect (*F*(3,40)=1.317, *p*=0.2823), meaning that ketamine’s impact depended on the timing of intervention and was not consistent across all timepoints.

**Figure 3.**
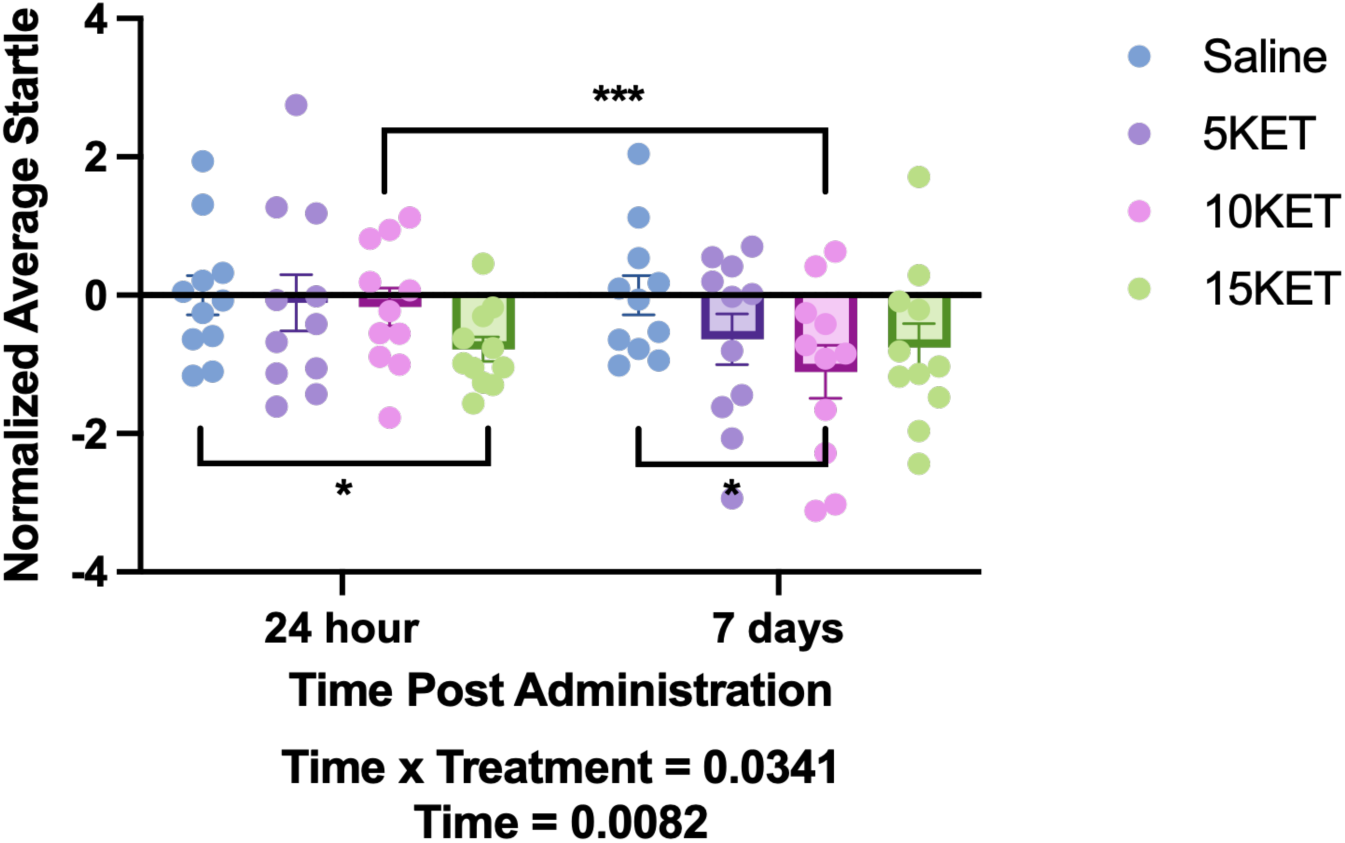
Effect of ketamine treatment on average startle response 24 hours (immediate trial) and 7 days (delayed trial) after ketamine administration by dose. +/- SEM. \**p*<0.05; \*\*\**p*<0.001.

Follow-up Šidák post hoc comparisons showed that despite most dose groups not differing significantly in average startle when comparing 24 hours to 7 day responses, rats treated with 10 mg/kg ketamine displayed a significant increase in average startle from 24 hours to 7 days (mean diff.=0.9389, Cl [0.2630, 1.615], *p*=0.0032), which indicates a delayed behavioral effect of the dose. No other doses nor saline vehicle controls showed significant changes between the two timepoints. Similarly, no significant pairwise comparisons emerged among treatment groups at either 24-hours or 7-day timepoints.

#### Normalized Average Startle by Sound Cue Intensity

A two-way ANOVA (treatment x cue intensity) was conducted to examine whether ketamine influenced responses across the three stimulus intensities of 95, 105, and 115 dB for both trials. Analyses revealed no significant immediate or delayed main effect of intensity (*F*(1.877, 73.21)=0.007750, *p*=0.9900; *F*(1.689, 67.56)=0.06063, *p*=0.9161) and no main effect of treatment for either timepoint (*F*(3, 40)=1.132, *p*=0.3477; *F*(3, 40)=1.471, *p*=0.2369), indicating that behavioral responses did not differ significantly across sound cue intensities or among ketamine groups on their own (Figure 4). Additionally, the analysis showed a trend-level interaction between intensity and treatment for the immediate trial (*F*(5.631, 73.21)=2.187, *p*=0.0576), suggesting the possibility of dose-dependent modulation across stimulus intensities. No interaction between intensity and treatment was recorded for the delayed trial (*F*(5.067, 67.56)=0.6073, *p*=0.6967).

**Figure 4.**
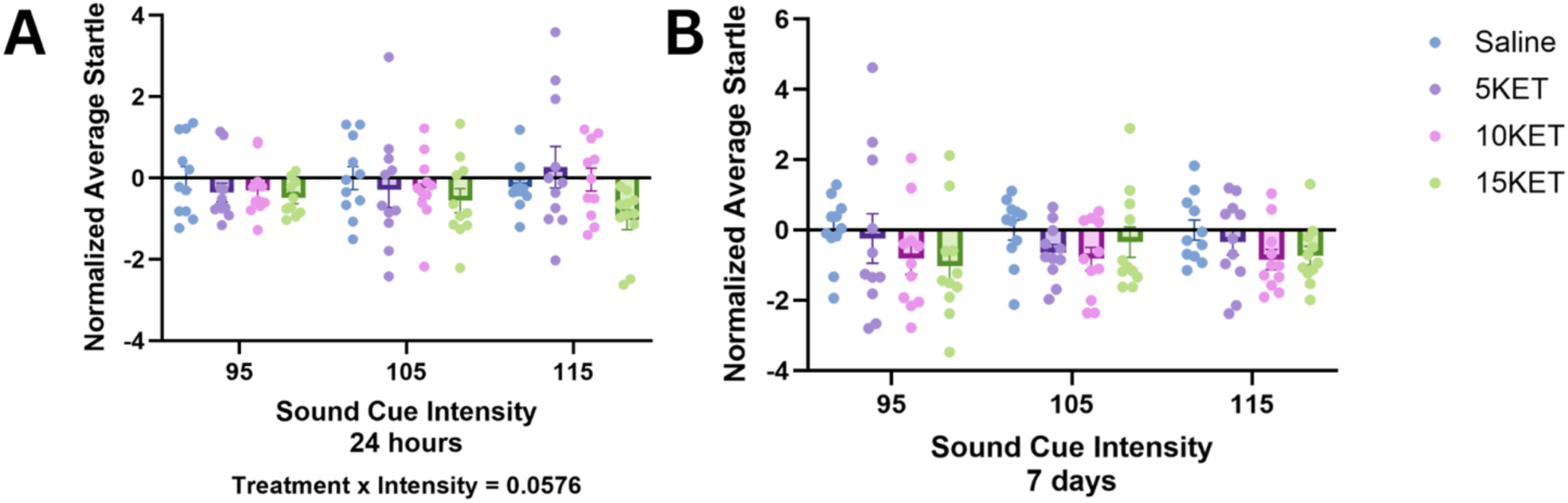
Trending interaction between ketamine treatment and sound cue intensity in average startle response. +/- SEM.

**Figure 5.**
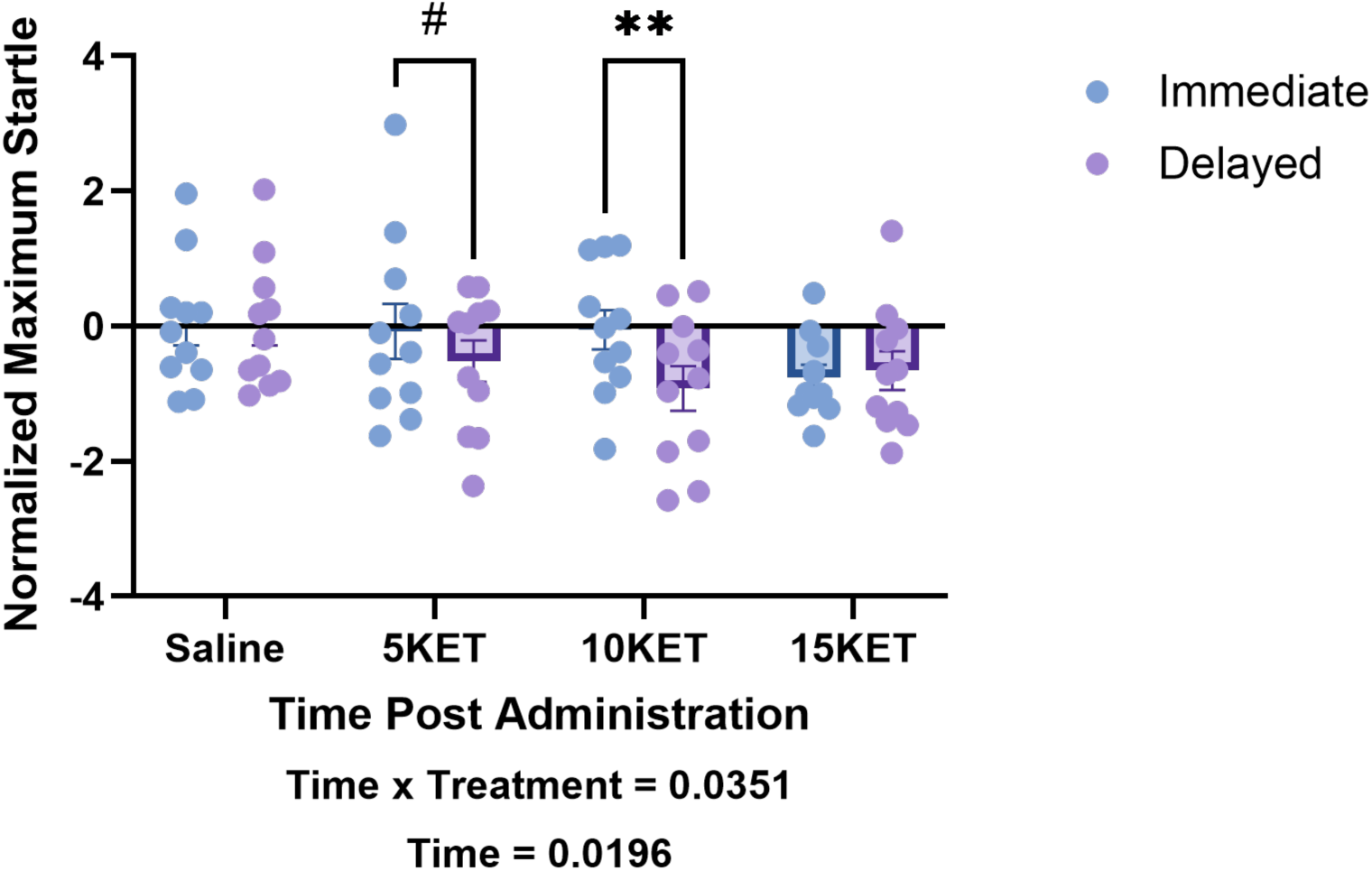
Effect of ketamine treatment on maximum startle response 24 hours (immediate trial) and 7 days (delayed trial) after ketamine administration by dose. +/- SEM. \**p*<0.05; \*\**p*<0.01; #*p*<0.08.

Follow-up Šidák multiple comparisons did not identify any statistically significant differences between treatment groups regardless of intensity level. Likewise, within each treatment condition, no significant pairwise differences emerged across the 95, 105, or 115 dB intensities.

#### Normalized Maximum Startle

A two-way repeated measures ANOVA was carried out to investigate the effect of the three doses of ketamine on maximum startle response across the immediate and delayed trials. A significant interaction between treatment condition and time was recorded (*F*(3, 40)=3.156, *p*=0.0351), indicating that responses to ketamine, as measured by the maximum startle response, have differed 24 hours and 7 days post administration. There was a statistically significant main effect of time alone as well (*F*(1, 40)=5.911, *p*=0.0196). No main effect of treatment was found (*F*(3, 40)=1.174, *p*=0.3316).

Follow-up Šidák post hoc comparisons demonstrated a trend between value of maximum startle after 24 hours and 7 days the administration of ketamine (mean diff.= 0.4407, Cl [-0.06214, 0.9435], *p*=0.0841). Additionally, there was significance for difference in responses to the 10 mg/kg between the two trials (mean diff.=0.8651, Cl [0.3623, 1.368], *p*=0.0012), once again highlighting the markedly strong difference in responses to the dose based on time that has passed since administration.

#### Normalized Maximum Startle by Sound Cue Intensity

When results for maximum startle were separated based on the intensity of the sound cue, a two-way repeated measures ANOVA revealed a statistically significant effect of intensity for immediate (*F*(1.724, 67.23)=229.0, *p*<0.0001) and delayed (*F*(1.662, 64.82)=170.0, *p*<0.0001) trials. An interaction between intensity and treatment for immediate but not delayed trial has been recorded as well (*F*(5.171, 67.23)=2.423, *p*=0.0426; *F*(4.986, 64.82)=0.8768, *p*=0.5015). No effect of treatment alone has been recorded 24 hours (*F*(3, 40)=1.267, *p*=0.2987) or 7 days (*F*(3, 40)=1.973, *p*=0.1335) following ketamine administration.

Follow-up Šidák post hoc comparisons situate the source of significance in within-treatment groups by cue intensity, further proving the success of the experimental setup. The analysis is omitted for brevity, but the results are visualized in Figure 6.

**Figure 6.**
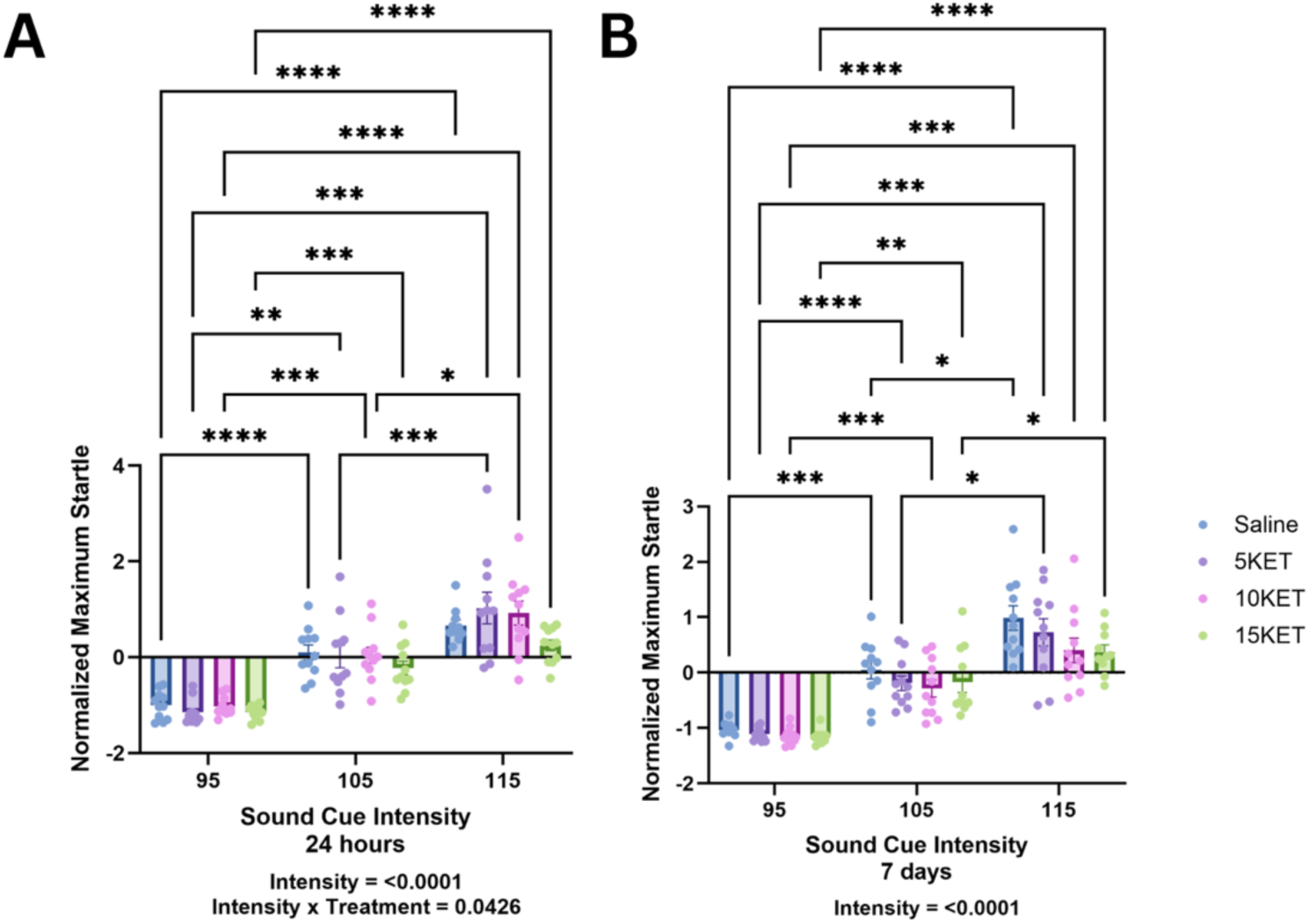
Interaction between ketamine treatment and sound cue intensity in maximum startle response. +/- SEM. \**p*<0.05; \*\**p*<0.01; \*\*\**p*<0.001; \*\*\*\**p*<0.0001.

### Immunofluorescence

#### 8-oxo-dG Intensity

To assess whether ketamine treatment had influence on global 8-oxo-dG intensity, a one-way ANOVA was performed for each of the four treatment groups for BLA, HPC (CA1, CA3, and HPC), PFC (IL and PL; Figure 7). The analysis revealed a significant main effect of treatment for BLA (*F*(3,24)=4.197, *p*=0.0160) and PL PFC (*F*(3,25)=3.308, *p*=0.0365; Figure 8). The findings indicate that ketamine doses significantly altered 8-oxo-dG global intensity values.

**Figure 7.**
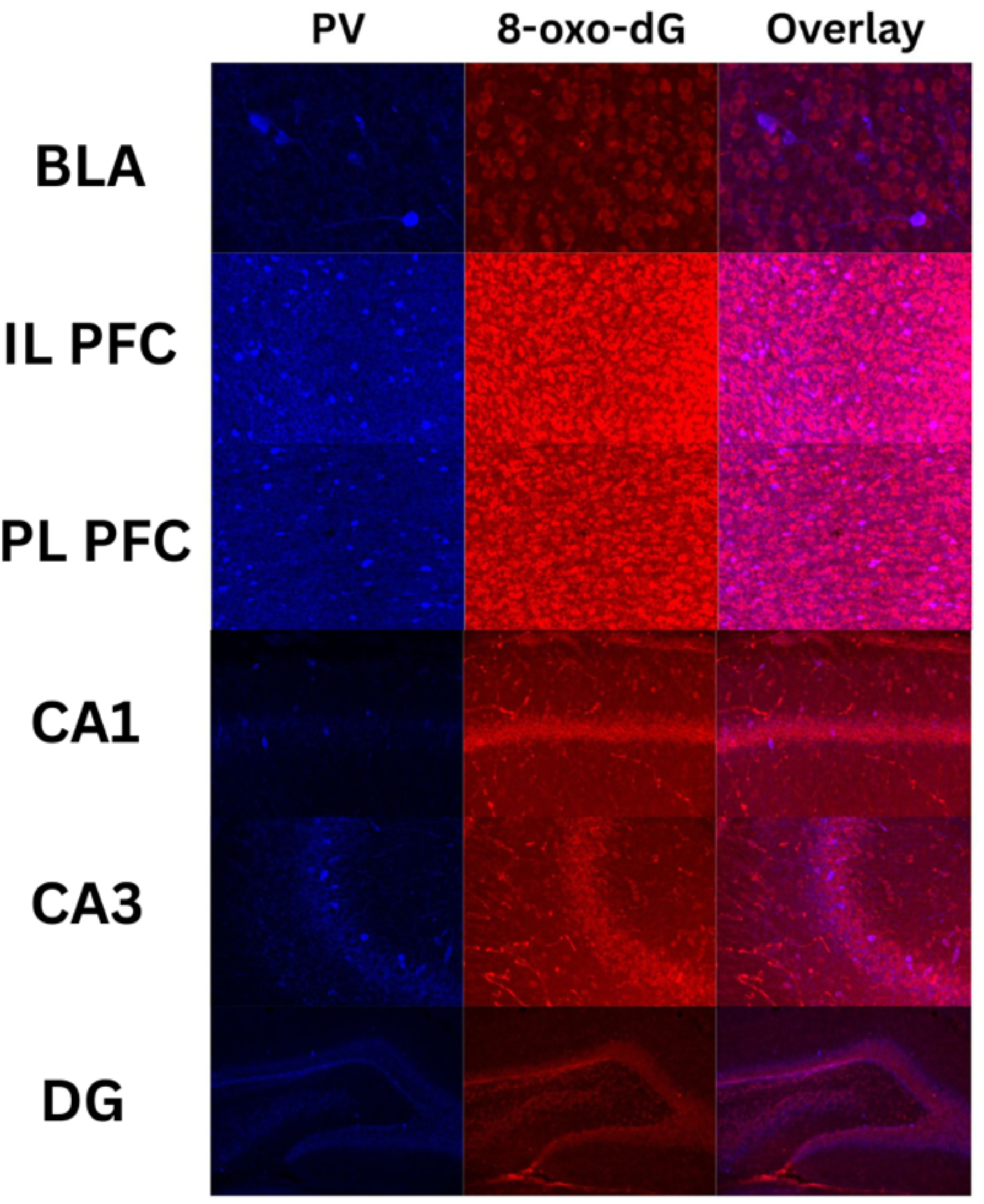
Representative images of immunohistochemistry staining for PV and 8-oxo-dG in BLA, PFC, and HPC.

**Figure 8.**
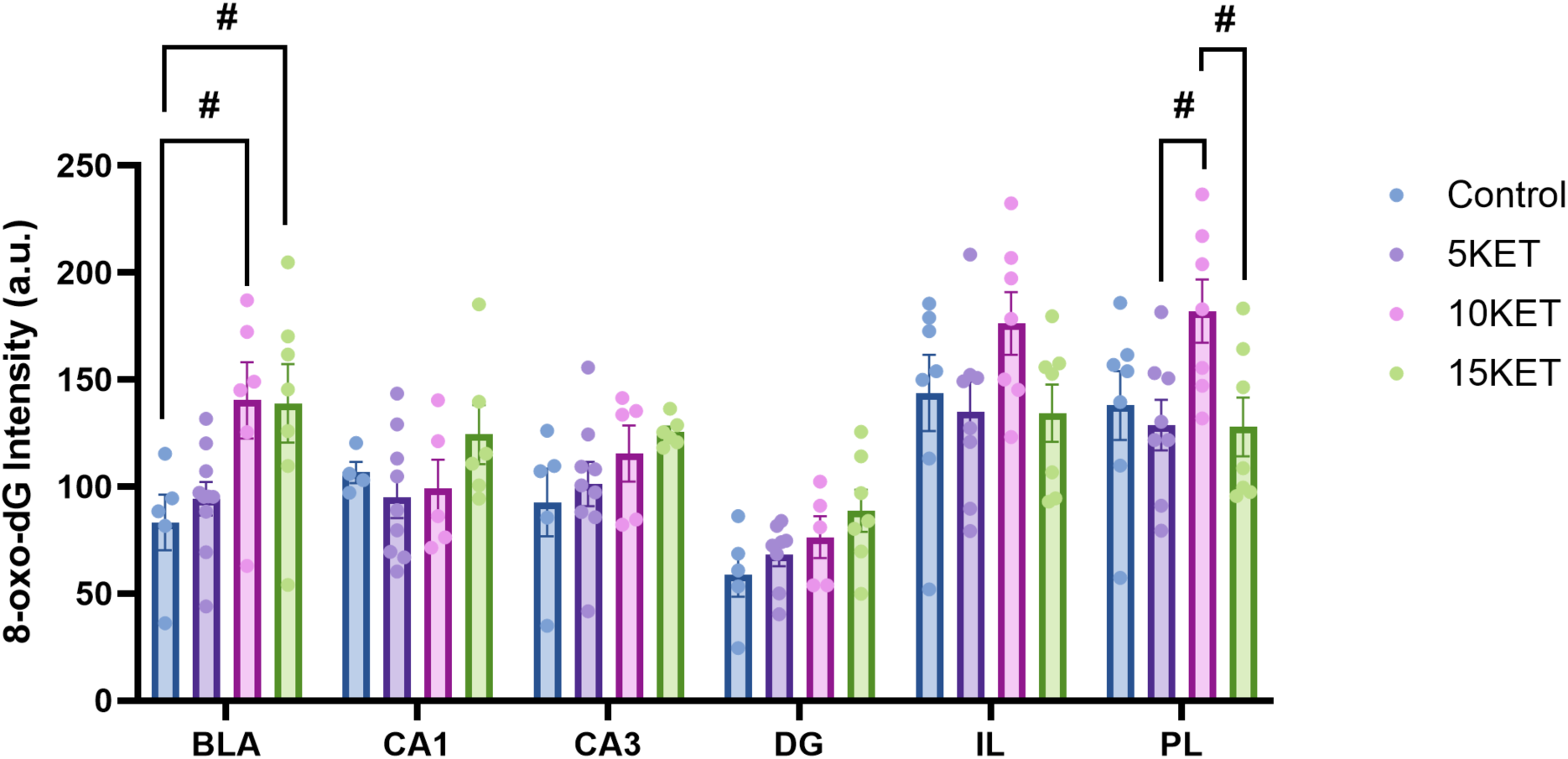
Global intensity of 8-oxo-dG intensity in the BLA, HPC (CA1, CA3, DG), and PFC (IL and PL) by treatment group. +/- SEM. #*p*<0.08; \**p*<0.05.

Follow-up Tukey post hoc comparisons showed that there was a trend for an increase in 8-oxo-dG intensity in the BLA between saline and 10 mg/kg (mean diff.= - 57.02, Cl [-118.4, 4.398], *p*=0.0755) and saline and 15 mg/kg (mean diff.=-55.54, Cl [-114.9, 3.848], *p*=0.0727) of ketamine. Additionally, a trend for an increase in oxidation was found in the PL PFC between 5 and 10 mg/kg (mean diff.= -53.41, Cl [-107.1, 0.2793], *p*=0.0516) and 10 and 15 mg/kg (mean diff.= 54.21, Cl [-1.237, 109.7], *p*=0.0570) of ketamine. No significant differences emerged within other brain regions, indicating that the effect was specific to the BLA and PL PFC.

#### PV Cell Number

A two-way repeated measures ANOVA (region x treatment) was conducted to evaluate differences in PV cell immunoreactivity across treatment groups for each brain region, which revealed no effect of treatment (*F*(3, 136)=1.252, *p*=0.2936), suggesting that ketamine did not significantly impact number of PV cells in any of the brain areas analyzed (Figure 9).

**Figure 9.**
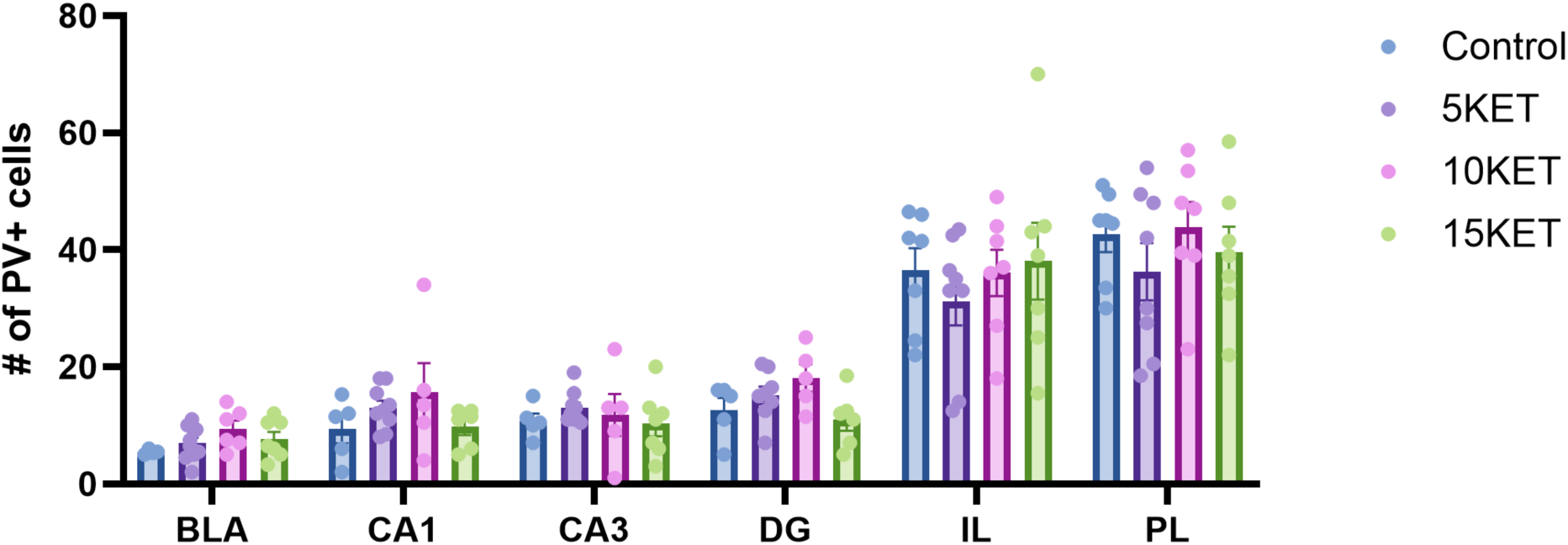
Number of PV+ cells in the BLA, HPC (CA1, CA3, DG), and PFC (IL and PL) by treatment group. +/- SEM.

#### 8-oxo-dG/PV Intensity

To determine whether ketamine treatment affected levels of oxidative damage specifically within PV+ interneurons, 8-oxo-dG signal intensity within PV+ cells was analyzed using a one-way ANOVA for each brain region. The analysis revealed a significant main effect of treatment for BLA (*F*(3, 23)=6.125, *p*=0.0032), indicating that ketamine dose affected the level of oxidative stress within the interneuron cells in the region (Figure 10).

**Figure 10.**
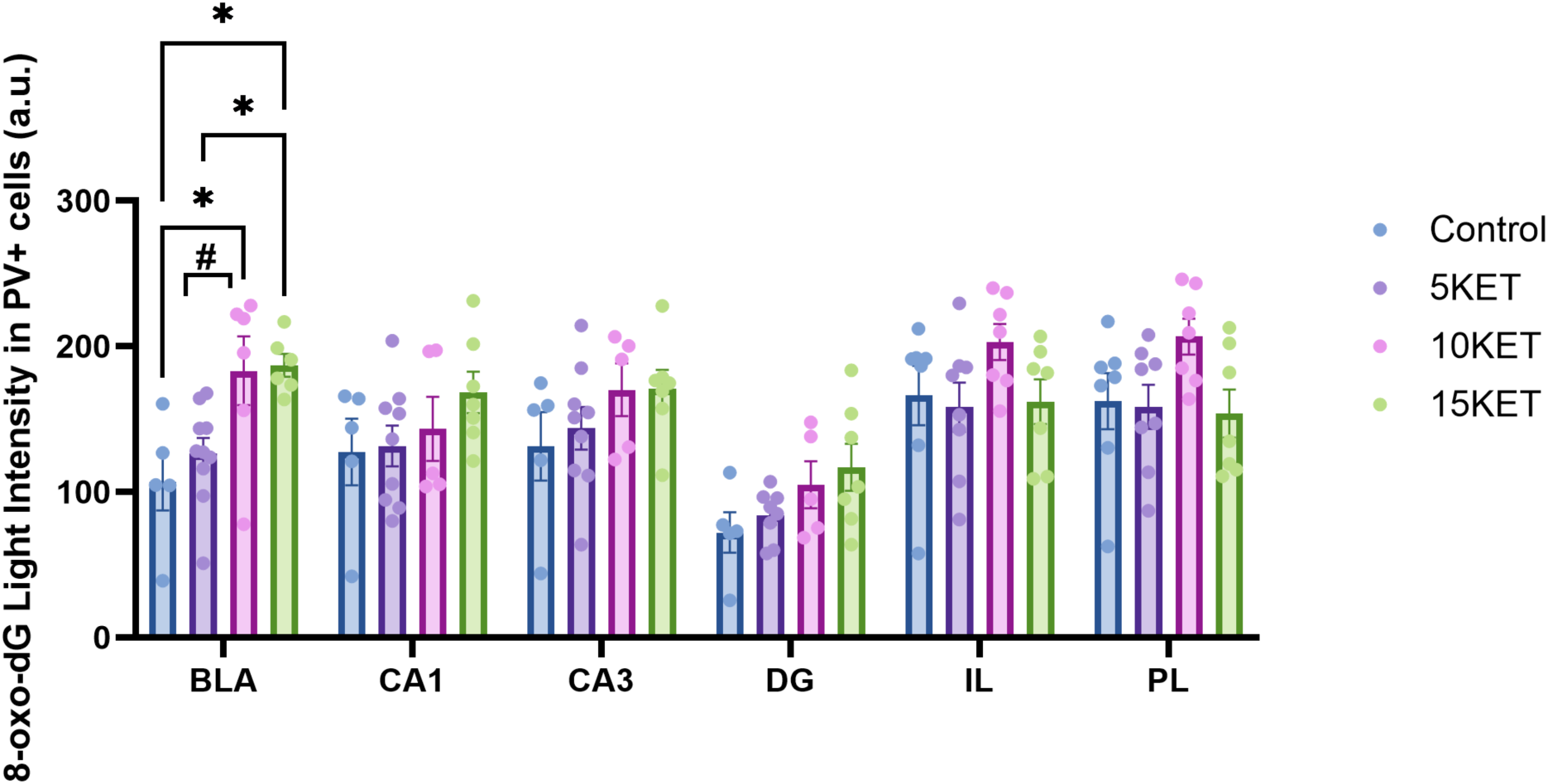
PV/8-oxo-dG colocalized intensity in the BLA, HPC (CA1, CA3, DG), and PFC (IL and PL) by treatment group. +/- SEM. \**p*<0.05; \*\**p*<0.01.

Tukey-corrected post hoc comparisons detected a significant increase in signal intensity in the BLA for saline and 10 mg/kg (mean diff.= -75.97, Cl [-143.1, -8.825], *p*= 0.0226), saline and 15 mg/kg (mean diff.= -79.78, Cl [-146.9, -12.64], *p*=0.0158), and 5 and 15 mg/kg (mean diff.= -60.57, Cl [-117.8, -3.312], *p*= 0.0355) of ketamine. Additionally, there was a trend for increase in signal intensity for 5 and 10 mg/kg (mean diff.= -56.76, Cl [-114.0, 0.5012], *p*= 0.0526) of ketamine. No other statistically significant differences for other brain regions were detected, which underlines a BLA-specific effect of ketamine on oxidative stress in PV+ cells.

## Discussion

The present study demonstrates that the effects of a single ketamine injection are time- and dose-dependent in female WKY rats. Across both average and maximum startle responses, the clearest therapeutic effect emerged at 10 mg/kg, marked by a significant reduction in startle response at 7 days compared to 24 hours. In comparison, the dose with the highest anxiolytic effect 24 hours post administration was 15 mg/kg, although it’s therapeutic benefits were not sustained at the 7-day test. As treatment alone did not produce a significant main effect across the three doses, the data suggest that ketamine’s behavioral impact depends more on when animals are tested after administration than on dose alone. Thus, the 10 mg/kg finding is most consistent with a delayed attenuation to hyperreactivity to stress-inducing cues rather than an immediate anxiolytic-like effect alone, which might better characterize the 15 mg/kg dose. Overall, this finding is consistent with existing literature on the role of ketamine as an anxiolytic, with multiple papers citing the clearest benefit for as long as 15 days after injection for the 10 mg/kg dose, with 15 mg/kg producing a mixed effect long-term (Lemeshova et al., 2026).

The results are furthermore consistent with the broader literature on ketamine as a therapeutic, suggesting that its effects are not always expressed as a simple reduction or amelioration of maladaptive stress response (Gass et al., 2019; Krystal et al., 2023; Sen et al., 2023). The outlines results suggest that ketamine might alter dynamics of reactivity to startle in WKY rats as a genetic model of treatment-resistant affective dysfunction, which once again aligns with existing literature citing greater responsiveness to therapeutic effects of ketamine within the strain (Manduca et al., 2020; Redei et al., 2023).

Considering that maximum startle increased strongly with louder pulses and that there was an immediate interaction between treatment condition and sound intensity, the results further suggest that ketamine might indirectly scale the degree of defensive response to more salient or pronounced sensory input even in the absence of strong treatment-group differences, exemplified by the decrease is average and maximum startle to 115 dB pulses in the 15 mg/kg ketamine condition. The demonstrated scaling is made more significant by the fact that acoustic startle is a highly translational paradigm with amygdala-centered circuitry that is often disrupted in affective disorders (Koch and Schnitzler, 1997; Ray et al., 2008; Seiglie et al., 2019).

Interestingly enough, to our knowledge, the current study is the first to demonstrate a clear anxiolytic effect of ketamine treatment in a female preclinical model of affective dysfunction, with previously only Fitzgerald et al. (2021) reporting a decrease in grooming behavior in the grooming test and increased center distance and center entrance frequency in the OFT 24 hours following the acute administration of 30 mg/kg i.p. ketamine in an unpredictable mild stress mouse C57BL/6J mouse model. In comparison, Radford et al. (2022) and Idil et al. (2025) found no anxiolytic effect of ketamine 24 hours or 2 days post administration in a Wistar and Sprague Dawley female model of stress, although the experimental paradigms utilized by the two studies were highly disparate and utilized different administration routes, dosing amount and timing, as well as the stress paradigm itself and behavioral measurement type. Thus, this study provides evidence for WKY rats as a reliable model of affective dysfunction for the study of ketamine as a novel antidepressant and anxiolytic, the use of which is extended to both male and female animals. Furthermore, the results suggest that an acute dose of 10 mg/kg i.p. ketamine might be sufficient to elicit a long-term anxiolytic effect within female WKY rats, while a higher dose of 15 mg/kg could confer a more immediate measure for reduction in anxiety-like behavior.

The immunohistochemical results argue against a simple interpretation of ketamine as neuroprotective and antioxidant, which is line with literature highlighting potential oxidation-promoting effects of ketamine, especially in the case of 8-oxo-dG as a marker of DNA damage globally and within PV+ cells specifically (Liu et al., 2012; Schiavone et al., 2019). Ketamine significantly increased global 8-oxo-dG intensity, with its most prominent effects emerging in the BLA and PL PFC. Further, ketamine treatment also increased oxidative stress signal intensity within PV+ cells specifically in the BLA. The number of PV+ cells, however, was not impacted by treatment in any of the investigated brain areas. Taken together, these findings suggest that acute ketamine exposure in this model altered the oxidative state of the stress circuitry without significantly changing the number of PV interneurons, which underscores the effect of ketamine as carrying an increased oxidative burden rather than reduced oxidative stress within the GABAergic PV interneurons in the BLA, as well as in unidentified cell type in the paralimbic area of the PFC.

The mechanism behind ketamine’s oxidative properties, however, might be more complicated than solely conferred neurological damage, as is evident from a clear therapeutic behavioral benefit emerging at the same doses of 10 and 15 mg/kg that resulted in highest expression of the marker within the PFC–BLA circuitry. While the current study did not detect change in number of PV interneurons, increased level of 8-oxo-dG marker within this subpopulation is consistent with previous reports in glutamate-GABA circuit disruption in rodents such as that of Schiavone et al. (2020), which discrepancy could potentially be explained in differences in dose and age of administration, as the aforementioned study used an acute dose of 30 mg/kg i.p. at PNDs 7, 9, and 11, corresponding to the neonatal life period in mice. Coupled with a demonstrated anxiolytic effect in behavior, it is possible to theorize that a ketamine-caused disruption to an already maladaptively formed PFC–BLA circuitry in a stress model could be beneficial at sub-toxic levels, leading to changes in glutamatergic and GABAergic signaling between the two regions.

Interestingly, one of proposed theories behind the pronounced affectively dysfunctional profile in WKY rats is a genetic deficiency in synaptic plasticity for projections between PL PFC and BLA, such as that lesions in inflicted in the area of Sprague Dawley rats resulted in a similar increase in extinction-resistant avoidance characteristic of the former strain (Fragale et al., 2017). As increases in reactive oxygen species (ROS), including 8-oxo-dG, can contribute to long-term potentiation, the lower subanesthetic doses used in this study might have been enough to allow for compensatory rewiring between the two brain regions without triggering cell death (Hahm et al., 2022). For comparison, a Wistar rat model of ketamine abuse that led to mass apoptotic cell death in unspecified cell populations of the cerebral cortex included 8 weeks of daily 50 mg/kg injections of ketamine, which is a markedly higher in dose than administered in this study (Sabziparvar et al., 2025).

In line with this balancing hypothesis, Liu et al. (2012) demonstrated an increase in NMDA receptor expression paired with an increase in ROS as measured by 8-oxo-dG production in neuronal rat tissue. According to the authors, neurons exposed to ketamine demonstrated elevated calcium influx and higher intracellular concentrations, which led to the compensatory NMDA upregulation (Liu et al., 2012). As PV neurons are highly sensitive to and dependent on calcium, a sudden increase in calcium flow would normally lead to rapid buffering to mitigate the increased input into the excitatory networks, such as that of pyramidal glutamatergic cells. Ketamine, however, blocks NMDA receptors on PV neurons, reducing PV firing and GABAergic inhibitory output, which likely results in more irregular excitatory glutamate firing and stronger dendritic calcium transients in pyramidal neurons, leading to increased expression in dependent plasticity pathways such as BDNF and mTOR (Aleksandrova et al., 2017; Zanos and Gould, 2018). This stronger activity recruits mitochondria, NADPH oxidases like NOX2, and calcium-sensitive metabolic enzymes to buffer the calcium influx, producing ROS in the process, which can be detected via 8-oxo-dG (Lisek et al., 2020).

Thus, when present at moderate levels, ROS may facilitate long-term potentiation by suppressing phosphatase activity via oxidation and inhibition and favoring kinase-driven synaptic strengthening (Knapp and Klann, 2002). In other words, our results suggest that when administered acutely at a low dose, ketamine might trigger the synaptic strengthening pathway described above, leading to circuit rewiring in circuits contributing to this model’s predisposition to depressive- and anxiety-like behaviors. The optimal window for such rewiring, however, is likely highly sensitive and narrow, resulting in cell death over therapeutic benefit, which might explain the decrease in anxiolytic effects long-term for the higher 15 mg/kg dose.

Overall, the present study shows that in female WKY rats, a single ketamine dose produced a delayed and dose-sensitive behavioral effect, while simultaneously increasing oxidative DNA markers in limbic and prefrontal regions. These findings suggest that ketamine’s actions are region-specific and time sensitive. Future work should determine whether the behavioral changes seen at 10 mg/kg reflect adaptive circuit remodeling or altered ketamine responsivity in females genetically predisposed to stress.

## Funding

This work was funded in part by Bowdoin College’s Grua/O’Connell Research Award (awarded to AL). Acoustic startle equipment and software was provided by the Faculty for Undergraduate Neuroscience Equipment Loan Program sponsored by San Diego Instruments (awarded to JAH).

## Notes

### Competing Interest Statement

The authors have declared no competing interest.

## References

1. Acevedo J, Mugarura NE, Welter AL, Johnson EM, Siegel JA. 2022. The effects of acute and repeated administration of ketamine on memory, behavior, and plasma corticosterone levels in female mice. Neuroscience. 512:99–109. doi:10.1016/j.neuroscience.2022.12.002. https://doi.org/10.1016/j.neuroscience.2022.12.002.

2. Ahmed GK, Elserogy YM, Elfadl GMA, Ghada Abdelsalam K, Ali MA. 2023. Antidepressant and anti-suicidal effects of ketamine in treatment-resistant depression associated with psychiatric and personality comorbidities: A double-blind randomized trial. Journal of Affective Disorders. 325:127–134. Doi: 10.1016/j.jad.2023.01.005.

3. Aleksandrova LR, Wang YT, Phillips AG. 2017. Hydroxynorketamine: Implications for the NMDA receptor hypothesis of ketamine’s antidepressant action. Chronic Stress. 1. doi:10.1177/2470547017743511. https://doi.org/10.1177/2470547017743511.

4. Almeida TM, Lacerda da Silva UR, Pires JP, Borges IN, Martins CRM, Cordeiro Q, Uchida RR (2021) Effectiveness of Ketamine for the Treatment of Post-Traumatic Stress Disorder – A Systematic Review and Meta-Analysis. Clin Neuropsychiatry 21:22–31. 10.36131/cnfioritieditore20240102

5. Arjmand S, Vadstrup Pedersen M, Silva NR, Landau AM, Joca S, Wegener G (2023) Sex and Estrous Cycle Are Not Mediators of S-Ketamine’s Rapid-Antidepressant Behavioral Effects in a Genetic Rat Model of Depression. Int J Neuropsychopharmacol 26:350–358. 10.1093/ijnp/pyad016

6. Binette AN, Liu J, Bayer H, Crayton KL, Melissari L, Sweck SO, Maren S. 2023. Parvalbumin-Positive interneurons in the medial prefrontal cortex regulate Stress-Induced fear extinction impairments in male and female rats. Journal of Neuroscience. 43(22):4162–4173. doi:10.1523/jneurosci.1442-22.2023. https://doi.org/10.1523/jneurosci.1442-22.2023.

7. Bogdańska-Chomczyk, E., Równiak, M., Huang, A. C., & Kozłowska, A. (2024). Parvalbumin interneuron deficiency in the prefrontal and motor cortices of spontaneously hypertensive rats: an attention-deficit hyperactivity disorder animal model insight. Frontiers in Psychiatry, 15, 1359237. 10.3389/fpsyt.2024.1359237

8. Bokma WA, Batelaan NM, Penninx BWJH, van Balkom AJLM (2021) Evaluating a dimensional approach to treatment resistance in anxiety disorders: A two-year follow-up study. Journal of Affective Disorders Reports 4:100139. 10.1016/j.jadr.2021.100139

9. Can A, Dao DT, Arad M, Terrillion CE, Piantadosi SC, Gould TD. 2011. The mouse forced swim test. Journal of Visualized Experiments.(58):e3638. doi:10.3791/3638. https://doi.org/10.3791/3638.

10. Dames S, Kryskow P, Watler C. 2022. A Cohort-Based Case Report: The Impact of Ketamine-Assisted therapy embedded in a Community of Practice Framework for healthcare providers with PTSD and Depression. Frontiers in Psychiatry. 12. doi:10.3389/fpsyt.2021.803279. https://doi.org/10.3389/fpsyt.2021.803279.

11. Deka B, Dash B, Bharali A, Ahmed A. 2022. Ketamine: More than Just NMDA Blocker. In: IntechOpen eBooks. 10.5772/intechopen.101113.

12. Drevets, W. C., Price, J. L., & Furey, M. L. (2008). Brain structural and functional abnormalities in mood disorders: implications for neurocircuitry models of depression. Brain Structure and Function, 213(1–2), 93–118. 10.1007/s00429-008-0189-x

13. Edwards, E., & King, J. (2009). Stress response: genetic consequences. In Elsevier eBooks (pp. 495–503). 10.1016/b978-008045046-9.00096-6

14. Elersič K, Banjac A, Živin M, Zorović M. 2025. Increased sensitivity to psychomotor effects of ketamine enantiomers in the Wistar-Kyoto depression model. Journal of Psychiatric Research. 184:307–317. doi:10.1016/j.jpsychires.2025.02.061. https://doi.org/10.1016/j.jpsychires.2025.02.061.

15. Gass N, Becker R, Reinwald J, Cosa-Linan A, Sack M, Weber-Fahr W, Vollmayr B, Sartorius A. 2019. Differences between ketamine’s short-term and long-term effects on brain circuitry in depression. Translational Psychiatry. 9(1):172. 10.1038/s41398-019-0506-6.

16. Gildawie, K. R., Honeycutt, J. A., & Brenhouse, H. C. (2019). Region-specific effects of maternal separation on perineuronal net and parvalbumin-expressing interneuron formation in male and female rats. Neuroscience, 428, 23–37. 10.1016/j.neuroscience.2019.12.010

17. Gürler, H. Ş., Bilgici, B., Akar, A. K., Tomak, L., & Bedir, A. (2014). Increased DNA oxidation (8-OHdG) and protein oxidation (AOPP) by low level electromagnetic field (2.45 GHz) in rat brain and protective effect of garlic. International Journal of Radiation Biology, 90(10), 892–896. 10.3109/09553002.2014.922717

18. Gutiérrez-Rojas, L., Porras-Segovia, A., Dunne, H., Andrade-González, N., & Cervilla, J. A. (2020). Prevalence and correlates of major depressive disorder: a systematic review. pmc.ncbi.nlm.nih.gov. 10.1590/1516-4446-2019-0650

19. Hahm JY, Park J, Jang E-S, Chi SW. 2022. 8-Oxoguanine: from oxidative damage to epigenetic and epitranscriptional modification. Experimental & Molecular Medicine. 54(10):1626–1642. doi:10.1038/s12276-022-00822-z. https://doi.org/10.1038/s12276-022-00822-z.

20. Hillhouse TM, Rice R, Porter JH. 2019. What role does the (2R,6R)-hydronorketamine metabolite play in the antidepressant-like and abuse-related effects of (R)-ketamine? British Journal of Pharmacology. 176(19):3886–3888. doi:10.1111/bph.14785. https://doi.org/10.1111/bph.14785.

21. Idil E, Yuksel B, Sen Z, Unal G (2025) Estrogen receptor alpha (ERα) partially modulates ketamine’s sustained anxiolytic effects without altering its antidepressant properties in female rats. Psychoneuroendocrinology 177:107455. 10.1016/j.psyneuen.2025.107455

22. Iqbal J, Huang G-D, Xue Y-X, Yang M, Jia X-J. 2023. The neural circuits and molecular mechanisms underlying fear dysregulation in posttraumatic stress disorder. Frontiers in Neuroscience. 17:1281401. doi:10.3389/fnins.2023.1281401. https://doi.org/10.3389/fnins.2023.1281401.

23. Ivanović A, Petrović J, Stanić D, Nedeljković J, Ilić M, Jukić MM, Pejušković B, Pešić V. 2025. Single subanesthetic dose of ketamine exerts antioxidant and antidepressive-like effect in ACTH-induced preclinical model of depression. Molecular and Cellular Neuroscience. 133:104006. doi:10.1016/j.mcn.2025.104006. https://doi.org/10.1016/j.mcn.2025.104006.

24. Jiang Y, Wang Y, Sun X, Lian B, Sun H, Wang G, Du Z, Li Q, Sun L (2017) Short- and long-term antidepressant effects of ketamine in a rat chronic unpredictable stress model. Brain and Behavior 7:e00749. 10.1002/brb3.749

25. Jiao X, Pang KCH, Beck KD, Minor TR, Servatius RJ. 2011. Avoidance perseveration during extinction training in Wistar-Kyoto rats: An interaction of innate vulnerability and stressor intensity. Behavioural Brain Research. 221(1):98–107. doi:10.1016/j.bbr.2011.02.029. https://doi.org/10.1016/j.bbr.2011.02.029.

26. Juneja, K., Afroze, S., Goti, Z., Sahu, S., Asawa, S., Bhuchakra, H. P., & Natarajan, B. (2024). Beyond therapeutic potential: a systematic investigation of ketamine misuse in patients with depressive disorders. Discover Mental Health, 4(1), 23. 10.1007/s44192-024-00077-2

27. Kessler RC, Petukhova M, Sampson NA, Zaslavsky AM, Wittchen H-U (2012) Twelve-month and lifetime prevalence and lifetime morbid risk of anxiety and mood disorders in the United States. Int J Methods Psychiatr Res 21:169–184. 10.1002/mpr.1359

28. Knapp LT, Klann E. 2002. Potentiation of hippocampal synaptic transmission by superoxide requires the oxidative activation of protein kinase C. Journal of Neuroscience. 22(3):674–683. doi:10.1523/jneurosci.22-03-00674.2002. https://doi.org/10.1523/jneurosci.22-03-00674.2002.

29. Koch, M., & Schnitzler, H. (1997). The acoustic startle response in rats—circuits mediating evocation, inhibition and potentiation. Behavioural Brain Research, 89(1–2), 35–49. 10.1016/s0166-4328(97)02296-1

30. Kraeuter A-K, Guest PC, Sarnyai Z. 2018. The Open Field Test for Measuring Locomotor Activity and Anxiety-Like Behavior. Methods in Molecular Biology. 1916:99–103. doi:10.1007/978-1-4939-8994-2_9. https://doi.org/10.1007/978-1-4939-8994-2_9.

31. Krystal JH, Kaye AP, Jefferson S, Girgenti MJ, Wilkinson ST, Sanacora G, Esterlis I. 2023. Ketamine and the neurobiology of depression: Toward next-generation rapid-acting antidepressant treatments. Proceedings of the National Academy of Sciences. 120(49):e2305772120. doi:10.1073/pnas.2305772120. https://doi.org/10.1073/pnas.2305772120

32. Lee Y, Davis M. 1997. Role of the hippocampus, the bed nucleus of the stria terminalis, and the amygdala in the excitatory effect of Corticotropin-Releasing hormone on the acoustic startle reflex. Journal of Neuroscience. 17(16):6434–6446. doi:10.1523/jneurosci.17-16-06434.1997. https://doi.org/10.1523/jneurosci.17-16-06434.1997.

33. Lehmann J. 1999. Sex differences in the acoustic startle response and prepulse inhibition in Wistar rats. Behavioural Brain Research. 104(1-2):113–117. 10.1016/s0166-4328(99)00058-3.

34. Lei, Y., & Tejani-Butt, S. (2010). N-methyl-d-aspartic acid receptors are altered by stress and alcohol in Wistar–Kyoto rat brain. Neuroscience, 169(1), 125–131. 10.1016/j.neuroscience.2010.05.003

35. Lemeshova A, Patel KA, Bloom AJ, Golan L, Haidari H, Limbada A, Zhao C, Honeycutt JA. 2025. A systematic review of ketamine’s anxiolytic potential in rodent behavioral models of anxiety and PTSD. Pharmacology Biochemistry and Behavior. 259:174144. doi:10.1016/j.pbb.2025.174144. https://doi.org/10.1016/j.pbb.2025.174144.

36. Li, Q., Zhao, W., Liu, S., Zhao, Y., Pan, W., Wang, X., Liu, Z., & Xu, Y. (2022). Partial resistance to citalopram in a Wistar–Kyoto rat model of depression: An evaluation using resting-state functional MRI and graph analysis. Journal of Psychiatric Research, 151, 242–251. 10.1016/j.jpsychires.2022.04.010

37. Liang J, Wu S, Xie W, He H. 2018. Ketamine ameliorates oxidative stress-induced apoptosis in experimental traumatic brain injury via the Nrf2 pathway. Drug Design Development and Therapy. Volume 12:845–853. doi:10.2147/dddt.s160046. https://doi.org/10.2147/dddt.s160046.

38. Lisek M, Zylinska L, Boczek T. 2020. Ketamine and Calcium Signaling—A crosstalk for neuronal physiology and pathology. International Journal of Molecular Sciences. 21(21):8410. doi:10.3390/ijms21218410. https://doi.org/10.3390/ijms21218410.

39. Liu, F., Patterson, T. A., Sadovova, N., Zhang, X., Liu, S., Zou, X., Hanig, J. P., Paule, M. G., Slikker, W., & Wang, C. (2012). Ketamine-Induced neuronal damage and altered N-Methyl-D-Aspartate receptor function in rat primary forebrain culture. Toxicological Sciences, 131(2), 548–557. 10.1093/toxsci/kfs296

40. López-Rubalcava, C. (2000). Strain differences in the behavioral effects of antidepressant drugs in the rat forced swimming test. Neuropsychopharmacology, 22(2), 191–199. 10.1016/s0893-133x(99)00100-1

41. Lu B, Nagappan G, Guan X, Nathan PJ, Wren P (2013) BDNF-based synaptic repair as a disease-modifying strategy for neurodegenerative diseases. Nat Rev Neurosci 14:401– 416. 10.1038/nrn3505

42. Manduca, J. D., Thériault, R., Williams, O. O., Rasmussen, D. J., & Perreault, M. L. (2020). Transient dose-dependent effects of ketamine on neural oscillatory activity in Wistar-Kyoto rats. Neuroscience, 441, 161–175. 10.1016/j.neuroscience.2020.05.012

43. McAuley JD, Stewart AL, Webber ES, Cromwell HC, Servatius RJ, Pang KCH. 2009. Wistar– Kyoto rats as an animal model of anxiety vulnerability: Support for a hypervigilance hypothesis. Behavioural Brain Research. 204(1):162–168. 10.1016/j.bbr.2009.05.036.

44. McDonnell, C. W., Dunphy-Doherty, F., Rouine, J., Bianchi, M., Upton, N., Sokolowska, E., & Prenderville, J. A. (2021). The Antidepressant-Like effects of a clinically relevant dose of ketamine are accompanied by biphasic alterations in working memory in the Wistar Kyoto Rat model of depression. Frontiers in Psychiatry, 11, 599588. 10.3389/fpsyt.2020.599588

45. McIntyre, R. S., Alsuwaidan, M., Baune, B. T., Berk, M., Demyttenaere, K., Goldberg, J. F., Gorwood, P., Ho, R., Kasper, S., Kennedy, S. H., Ly-Uson, J., Mansur, R. B., McAllister-Williams, R. H., Murrough, J. W., Nemeroff, C. B., Nierenberg, A. A., Rosenblat, J. D., Sanacora, G., Schatzberg, A. F., . . . Maj, M. (2023). Treatment-resistant depression: definition, prevalence, detection, management, and investigational interventions. World Psychiatry, 22(3), 394–412. 10.1002/wps.21120

46. Miller EA, Kastner DB, Grzybowski MN, Dwinell MR, Geurts AM, Frank LM. 2020. Robust and replicable measurement for prepulse inhibition of the acoustic startle response. Molecular Psychiatry. 26(6):1909–1927. doi:10.1038/s41380-020-0703-y. https://doi.org/10.1038/s41380-020-0703-y.

47. Morgan, C., Grillon, C., Southwick, S. M., Davis, M., & Charney, D. S. (1995). Fear-potentiated startle in posttraumatic stress disorder. Biological Psychiatry, 38(6), 378–385. 10.1016/0006-3223(94)00321-s

48. Murrough JW, Iosifescu DV, Chang LC, Al Jurdi RK, Green CE, Perez AM, Iqbal S, Pillemer S, Foulkes A, Shah A, et al. 2013. Antidepressant Efficacy of Ketamine in Treatment-Resistant Major Depression: A Two-Site Randomized Controlled Trial. American Journal of Psychiatry. 170(10):1134–1142. 10.1176/appi.ajp.2013.13030392.

49. Palmer, A., & Printz, M. (1999). Strain differences in Fos expression following airpuff startle in Spontaneously Hypertensive and Wistar Kyoto rats. Neuroscience, 89(3), 965–978. 10.1016/s0306-4522(98)00333-9

50. Perlman, G., Tanti, A., & Mechawar, N. (2021). Parvalbumin interneuron alterations in stress-related mood disorders: A systematic review. Neurobiology of Stress, 15, 100380. 10.1016/j.ynstr.2021.100380

51. Picard, N., Takesian, A. E., Fagiolini, M., & Hensch, T. K. (2019). NMDA 2A receptors in parvalbumin cells mediate sex-specific rapid ketamine response on cortical activity. Molecular Psychiatry, 24(6), 828–838. 10.1038/s41380-018-0341-9

52. Ponton E, Turecki G, Nagy C (2022) Sex Differences in the Behavioral, Molecular, and Structural Effects of Ketamine Treatment in Depression. Int J Neuropsychopharmacol 25:75–84. 10.1093/ijnp/pyab082

53. Primo MJ, Fonseca-Rodrigues D, Almeida A, Teixeira PM, Pinto-Ribeiro F. 2023. Sucrose preference test: A systematic review of protocols for the assessment of anhedonia in rodents. European Neuropsychopharmacology. 77:80–92. doi:10.1016/j.euroneuro.2023.08.496. https://doi.org/10.1016/j.euroneuro.2023.08.496.

54. Radford KD, Spencer HF, Zhang M, Berman RY, Girasek QL, Choi KH. 2019. Association between intravenous ketamine-induced stress hormone levels and long-term fear memory renewal in Sprague-Dawley rats. Behavioural Brain Research. 378:112259. doi:10.1016/j.bbr.2019.112259. https://doi.org/10.1016/j.bbr.2019.112259.

55. Ray, W. J., Molnar, C., Aikins, D., Yamasaki, A., Newman, M. G., Castonguay, L., & Borkovec, T. D. (2008). Startle response in Generalized Anxiety Disorder. Depression and Anxiety, 26(2), 147–154. 10.1002/da.20479

56. Redei, E. E., Udell, M. E., Woods, L. C. S., & Chen, H. (2022). The Wistar Kyoto Rat: A model of Depression Traits. Current Neuropharmacology, 21(9), 1884–1905. 10.2174/1570159x21666221129120902

57. Réus, G. Z., Carlessi, A. S., Titus, S. E., Abelaira, H. M., Ignácio, Z. M., Da Luz, J. R., Matias, B. I., Bruchchen, L., Florentino, D., Vieira, A., Petronilho, F., & Quevedo, J. (2015). A single dose of S-ketamine induces long-term antidepressant effects and decreases oxidative stress in adulthood rats following maternal deprivation. Developmental Neurobiology, 75(11), 1268–1281. 10.1002/dneu.22283

58. Ryazantseva M, Liiwand M, Shteinikov V, Kunnari A-J, Sulku J, Lauri SE. 2025. Parvalbumin interneurons gate amygdala excitability and response to chronic stress via kainate receptor-driven tonic GABAB receptor-mediated inhibition. Molecular Psychiatry. 30(11):5093–5107. doi:10.1038/s41380-025-03093-y. https://doi.org/10.1038/s41380-025-03093-y.

59. Sabbagh JJ, Murtishaw AS, Bolton MM, Heaney CF, Langhardt M, Kinney JW. 2013. Chronic ketamine produces altered distribution of parvalbumin-positive cells in the hippocampus of adult rats. Neuroscience Letters. 550:69–74. doi:10.1016/j.neulet.2013.06.040. https://doi.org/10.1016/j.neulet.2013.06.040.

60. Schiavone, S., Morgese, M. G., Bove, M., Colia, A. L., Maffione, A. B., Tucci, P., Trabace, L., & Cuomo, V. (2019). Ketamine administration induces early and persistent neurochemical imbalance and altered NADPH oxidase in mice. Progress in Neuro-Psychopharmacology and Biological Psychiatry, 96, 109750. 10.1016/j.pnpbp.2019.109750

61. Seiglie, M. P., Huang, L., Cottone, P., & Sabino, V. (2019). Role of the PACAP system of the extended amygdala in the acoustic startle response in rats. Neuropharmacology, 160, 107761. 10.1016/j.neuropharm.2019.107761

62. Sen ZD, Chand T, Danyeli LV, Kumar VJ, Colic L, Li M, Yemisken M, Javaheripour N, Refisch A, Opel N, et al. 2023a. The effect of ketamine on affective modulation of the startle reflex and its resting-state brain correlates. Scientific Reports. 13(1):13323. doi:10.1038/s41598-023-40099-4. https://doi.org/10.1038/s41598-023-40099-4

63. Serafini, G., Howland, R., Rovedi, F., Girardi, P., & Amore, M. (2014). The Role of Ketamine in Treatment-Resistant Depression: A Systematic Review. Current Neuropharmacology, 12(5), 444–461. 10.2174/1570159x12666140619204251

64. Singh, I., Morgan, C., Curran, V., Nutt, D., Schlag, A., & McShane, R. (2017). Ketamine treatment for depression: opportunities for clinical innovation and ethical foresight. The Lancet Psychiatry, 4(5), 419–426. 10.1016/s2215-0366(17)30102-5

65. Su, X., Dong, X., Lu, C., Zhang, M., Li, Y., Xiao, H., Wang, J., Sun, Y., Cong, B., & Wang, S. (2025). The mechanism of parvalbumin interneurons regulating glutamatergic neurons involvement in stress induced anxiety in the basolateral amygdala of male mice. Scientific Reports, 15(1), 26424. 10.1038/s41598-025-10130-x

66. Thelen C, Sens J, Mauch J, Pandit R, Pitychoutis PM. 2016. Repeated ketamine treatment induces sex-specific behavioral and neurochemical effects in mice. Behavioural Brain Research. 312:305–312. doi:10.1016/j.bbr.2016.06.041. https://doi.org/10.1016/j.bbr.2016.06.041.

67. Vaidyanathan U, Welo EJ, Malone SM, Burwell SJ, Iacono WG. 2013. The effects of recurrent episodes of depression on startle responses. Psychophysiology. 51(1):103–109. 10.1111/psyp.12152.

68. Will CC, Aird F, Redei EE. 2003. Selectively bred Wistar–Kyoto rats: an animal model of depression and hyper-responsiveness to antidepressants. Molecular Psychiatry. 8(11):925– 932. 10.1038/sj.mp.4001345.

69. Willner P. 1997. Validity, reliability and utility of the chronic mild stress model of depression: a 10-year review and evaluation. Psychopharmacology. 134(4):319–329. doi:10.1007/s002130050456. https://doi.org/10.1007/s002130050456.

70. Wright KN, Strong CE, Addonizio MN, Brownstein NC, Kabbaj M. 2016. Reinforcing properties of an intermittent, low dose of ketamine in rats: effects of sex and cycle. Psychopharmacology. 234(3):393–401. doi:10.1007/s00213-016-4470-z. https://doi.org/10.1007/s00213-016-4470-z.

71. Xiao, M., Zhong, H., Xia, L., Tao, Y., & Yin, H. (2017). Pathophysiology of mitochondrial lipid oxidation: Role of 4-hydroxynonenal (4-HNE) and other bioactive lipids in mitochondria. Free Radical Biology and Medicine, 111, 316–327. 10.1016/j.freeradbiomed.2017.04.363

72. Yang J, Wang N, Yang C, Shi J, Yu H, Hashimoto K. 2015. Serum Interleukin-6 Is a Predictive Biomarker for Ketamine’s Antidepressant Effect in Treatment-Resistant Patients With Major Depression. Biological Psychiatry. 77(3):e19–e20. 10.1016/j.biopsych.2014.06.021.

73. Yang X, Chen D (2024) Comparing the adverse effects of ketamine and esketamine between genders using FAERS data. Front Pharmacol 15. 10.3389/fphar.2024.132943

74. Yau JO-Y, Chaichim C, Power JM, McNally GP. 2021. The roles of basolateral amygdala parvalbumin neurons in fear learning. Journal of Neuroscience. 41(44):9223–9234. doi:10.1523/jneurosci.2461-20.2021. https://doi.org/10.1523/jneurosci.2461-20.2021.

75. Yavi, M., Lee, H., Henter, I. D., Park, L. T., & Zarate, C. A. (2022). Ketamine treatment for depression: a review. Discover Mental Health, 2(1), 9. 10.1007/s44192-022-00012-3

76. Zanos P, Gould TD. 2018. Mechanisms of ketamine action as an antidepressant. Molecular Psychiatry. 23(4):801–811. doi:10.1038/mp.2017.255. https://doi.org/10.1038/mp.2017.255.

